# Tumors widely express hundreds of embryonic germline genes

**DOI:** 10.1101/2020.09.08.287284

**Authors:** Jan Willem Bruggeman, Naoko Irie, Paul Lodder, Ans M.M. van Pelt, Jan Koster, Geert Hamer

## Abstract

We have recently described a class of 756 genes that are widely expressed in cancer, while normally restricted to adult germ cells, referred to as germ cell cancer genes (GC-genes). We hypothesized that carcinogenesis involves reactivation of biomolecular processes and regulatory mechanisms that, under normal circumstances, are restricted to germline development. This would imply that cancer cells share gene expression profiles with primordial germ cells (PGCs). We therefore compared the transcriptomes of human PGCs (hPGCs) and PGC-like cells (PGCLCs) with 17 382 samples from 54 healthy somatic tissues (GTEx) and 11 003 samples from 33 tumor types (TCGA), and identified 672 GC-genes, expanding the known GC-gene pool by 387 genes (51%). Because GC-genes specific to the embryonic germline are not expressed in any adult tissue, targeting these in cancer treatment may result in fewer side effects than targeting conventional cancer/testis (CT) or GC-genes and may preserve fertility. We anticipate that our extended GC-dataset enables improved understanding of tumor development and may provide multiple novel targets for cancer treatment development.

## Introduction

Many genes have been identified that drive the transition from healthy cells into cancer cells. Such oncogenes contribute to the acquisition of cancer-specific hallmarks^1,2^, such as uncontrolled cell divisions, angiogenesis, aberrant apoptosis regulation and telomere maintenance. Targeting these hallmark processes is effectively used by many current cancer therapies. However, because the majority of these processes are also widely used by non-cancerous cells, these therapies often cause severe side effects. To diminish potential side effects in cancer therapy, the identification of oncogenes that are inactive in mature healthy human tissues is paramount to the development of novel diagnostics and therapeutics.

One group of genes that has been studied to this end are cancer/testis (CT) genes^3,4^. CT-genes have been identified by selecting genes that are highly expressed in testis tissue and cancer, and expressed in a limited number of healthy somatic tissues. This approach has resulted in the identification of 1 128 CT-genes to date (**figure 1A**)^5,6^. However, CT-genes include the genes expressed in somatic cells in the testis, precluding the detection of true germ cell specific genes. By using the transcriptome of isolated adult male germ cells, we have recently identified 756 true germ cell specific genes expressed in cancer, termed germ cell cancer genes (GC-genes), of which 630 (83%) were newly identified (**figure 1A & 1B**)^7^.

**Figure 1.**
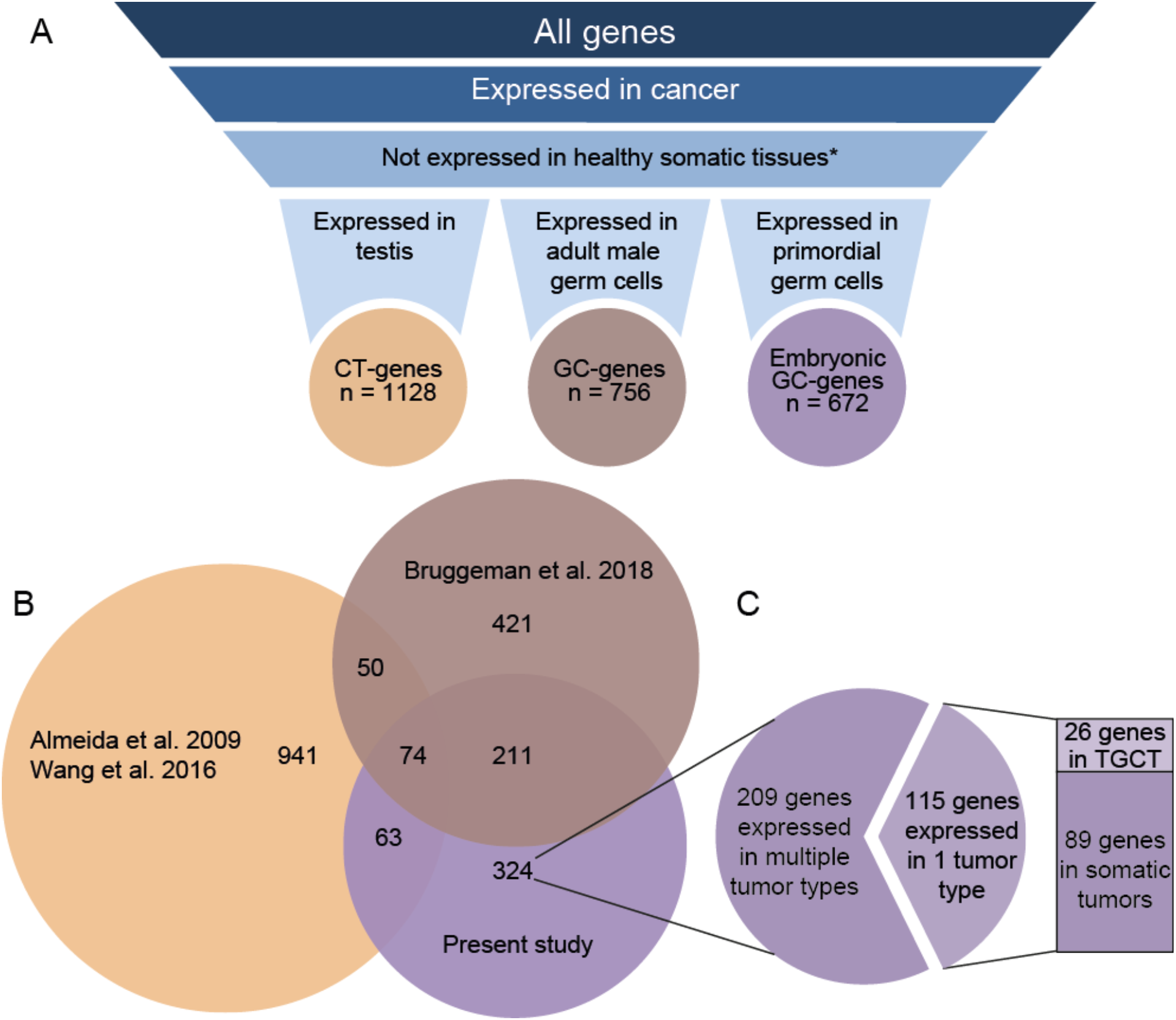
[a] Visualization of how CT and GC-genes have been identified. Left: genes included by Wang et al.^6^ and Almeida et al.^5^ have been based on gene expression in whole testis tissue (CT-genes). Middle: genes included by Bruggeman et al.^7^ have been based on gene expression in germ cells (GC-genes). Right: Genes included in the present analysis have been included based on gene expression in human primordial germ cells (GC-genes). * = testis and ovary were excluded as they are not considered somatic. **[b] Approximately half of the embryonic GC-genes have not yet been described before as GC-gene or CT-gene**. Venn diagram comparing the present analysis of human germline – cancer (GC) genes expressed in primordial germ cells (red) to earlier identified GC-genes expressed in adult germ cells^17^ and cancer/testis (CT) genes^5,6^ (**supplementary data 2C**). **[c] The majority of newly identified embryonic GC-genes are expressed in multiple tumor types**. From the 115 genes expressed in only one tumor type, 26 are expressed in testicular germ cell tumors (TGCT), showing that the majority of GC-genes are expressed in tumors that originate from somatic tissues.

Cancer and germ cells share many biological properties, such as “endless/immortal” propagation and developmental potential. Indeed, we found that GC-genes are involved in processes that can drive and maintain tumor development^7^. We now hypothesize that also cancer initiation and early development (carcinogenesis) involves biomolecular processes and regulatory mechanisms that are usually restricted to germ cell development. To investigate this, we sought the oncogenic potential of molecular regulation for the primordial germ cells (PGCs), the embryonic precursors of the adult germ cells, which has not been investigated. Human PGCs are specified in the week 2 embryo by expression of the transcription factor SOX17^8^, resulting in global hypomethylation and latent pluripotency. Moreover, the process can be recapitulated *in vitro* by using human pluripotent stem cells^8–10^. Specified PGCs in embryos multiply extensively and migrate from their site origin in the proximal epiblast, through the developing hindgut, to the gonadal ridges during week 3 to 5 of human development^11^. As such, the physiology of PGCs includes processes that also have been proposed as hallmarks of cancer, including continuous replicative potential via telomere lengthening^12^, deregulating cellular energetics^13^, as well as invasive potential and metastasis^14^. Furthermore, DNA hypomethylation, in itself a characteristic feature of PGCs^15^, is also a proposed consequence of germ cell specific gene activity in tumors^16^. As these processes favor development and survival of the cancer cell, investigating genes specific to PGCs and cancer has large potential to tumor biology. When a therapeutic target is unique to cancer and adult germ cells, side effects of targeting such gene products during cancer therapy would be limited to (temporary) infertility. However, targeting genes specific to PGCs may not even result in any side effects because these gene products are absent from the adult germ cells.

## Methods

### Used datasets

In order to be able to make a selection, we merged several publicly accessible RNA expression datasets (transcriptomes) into one file (**Supplementary data 1A**). In this process, we have used the transcriptomes of hPGCs of week 5.5 – 8.5 of human embryonic development and PGCLCs representing PGCs in week 2-3 of human embryonic development^8^ as base files. We then matched each gene. to the expression data from the following transcriptomes in one file (**Supplementary data 1A**):

1. The transcriptome of ESCs cultured in conventional media^8^
2. The transcriptome of embryonic somatic tissue, which surrounds the hPGCs in-situ^8^
3. The transcriptome of adult male germ cells in various stages of spermatogenesis^17^
4. The Genotype Tissue Expression (GTEx) project, containing 17 382 samples from 54 healthy (i.e. non-cancerous) tissues (**Supplementary data 1B**)^18^
5. The Cancer Genome Atlas (TCGA) project, containing 11 003 samples from 33 tumor types (**Supplementary data 1C**)^19^

Genes that had no expression data available from the GTEx and/or TCGA project were excluded (n = 1 728, **Supplementary data 1D**), leaving 15 992 genes for our analysis (**Supplementary data 1A**). From the GTEx database, we excluded 2 transformed cell lines and converted the data to a log_2_ scale. Because adult germ cells are present in testis and ovary tissue, we excluded “testis” and “ovary” as well in order to allow for the inclusion of previously identified CT and/or GC-genes. The RNA expression of both PGC types (hPGCs & PGCLCs), somatic gonadal tissue and ESCs was corrected for gene length (expression/length*1000) to reduce the number of false positives. The gene expression values in the other datasets had already been corrected for gene length.

*Supplementary data available on request; g*.*hamer@amsterdamumc*.*nl*

### Selection of genes

Because we compare gene expression levels from multiple sources with distinct distributions, we cannot simply compare these values between datasets. Thus, we determined a cut-off for each dataset to in- or exclude genes (**Supplementary figure 1**). To be sure that genes are expressed highly in hPGCs, the minimum expression of each gene in any stage (week 5.5 – 8.5) was compared between female and male hPGCs, after which the maximum value of the two was used to determine arbitrary inclusion criteria. For PGCLCs, inclusion criteria were based on the average of two samples. Genes showing hPGC expression >0.72 (33th percentile) or PGCLC expression >0.50 (33th percentile) were considered expressed (**Supplementary figure 1A**). Additional inclusion criteria were applied to the maximum RNA expression levels per gene in all tissue/tumor types. Namely, genes that are not expressed in any normal tissue (GTEx < 3.0, **Supplementary figure 1B**) and are expressed in at least one tumor type (TCGA > 2.3, **Supplementary figure 1C**) have been included. The values represent the average normalized log_2_(reads per million) in a varying number of patient samples (**Supplementary data 1B & 1C**). These criteria include genes in the following percentiles: 74% for hPGCs and PGCs, 13% for normal tissues and 89% for cancer. Because the inclusion criteria are arbitrary, we have developed a web-based application that allows anyone to manually change the inclusion criteria and observe how this affects the results: http://venn.lodder.dev or later: https://www.amsterdamresearch.org/web/reproduction-and-development/tools.htm.

### Data analysis

Gene ontology (GO) analysis was performed with DAVID Bioinformatics Resources^20^ v6.8. Cellular component analysis was performed by using the Panther 10.0 classification system^21^. Genes associated with the cell surface were those attributed to GO-term 9986. Data visualization was done in R2^22^ and the JavaScript library D3. Protein expression was evaluated using the Human Protein Atlas (HPA)^23^, available from http://www.proteinatlas.org. Only proteins categorized as “Not detected” in all tissues except the seminiferous tubules and ovaries were classified as “validated”. The GC-signature scores were attributed using the ‘sample ranked geneset scores’ function in R2, which takes a list of genes and ranks these genes based on expression in a provided set of samples, such as each of the 917 cell lines in the CCLE^24^ or each of 515 lung adenocarcinoma tumor samples in the TCGA dataset^19^. The signature score is the average percentile of these ranks, and may thus be used as a measure for a cancer cell line’s similarity to the germline.

## Results

To assemble an inventory of genes of interest, we explored gene expression of tumor data from the TCGA^19^, normal tissue expression from GTEx^18^, and primordial germ cell data from Irie et al. 2015^8^. By applying similarly strict inclusion criteria as in our previous study^7^ (**Supplementary figure 1**), we here identify 672 genes that are expressed in primordial germ cells (i.e. either human PGCs derived from week 5.5 – 8.5 embryos and/or *in vitro* derived PGCLCs representing week 2.5 – 3.0 of development) and a wide variety of tumor types, while being virtually undetectable from the GTEx database of healthy somatic tissues (**Figure 1A, Supplementary data 2A**). Because they are expressed in the embryonic germline, we will refer to these genes as “embryonic” GC-genes. 348 genes (51%) are also expressed in mature germ cells or testis tissue and have been identified as GC or CT-genes before (**Figure 1B, Supplementary data 2B**). We thus expand the known CT/GC-gene pool^5–7^ with 324 new genes that are restricted to the germline and cancer (**Figure 1C**). We have visualized how custom inclusion criteria affect the results and their overlap with other studies in a web-based application, available from http://venn.lodder.dev or later: www.amsterdamresearch.org/web/reproduction-and-development/tools.htm. A gene ontology (GO) analysis suggests that the 672 genes expressed in PGCs and cancer cells play a role in unique processes, including the meiotic cell cycle, nucleic acid metabolic processes, nuclear division, strand displacement, gene regulation, and stem cell population maintenance (**Table 1, Supplementary data 2C**).

**Table 1.**
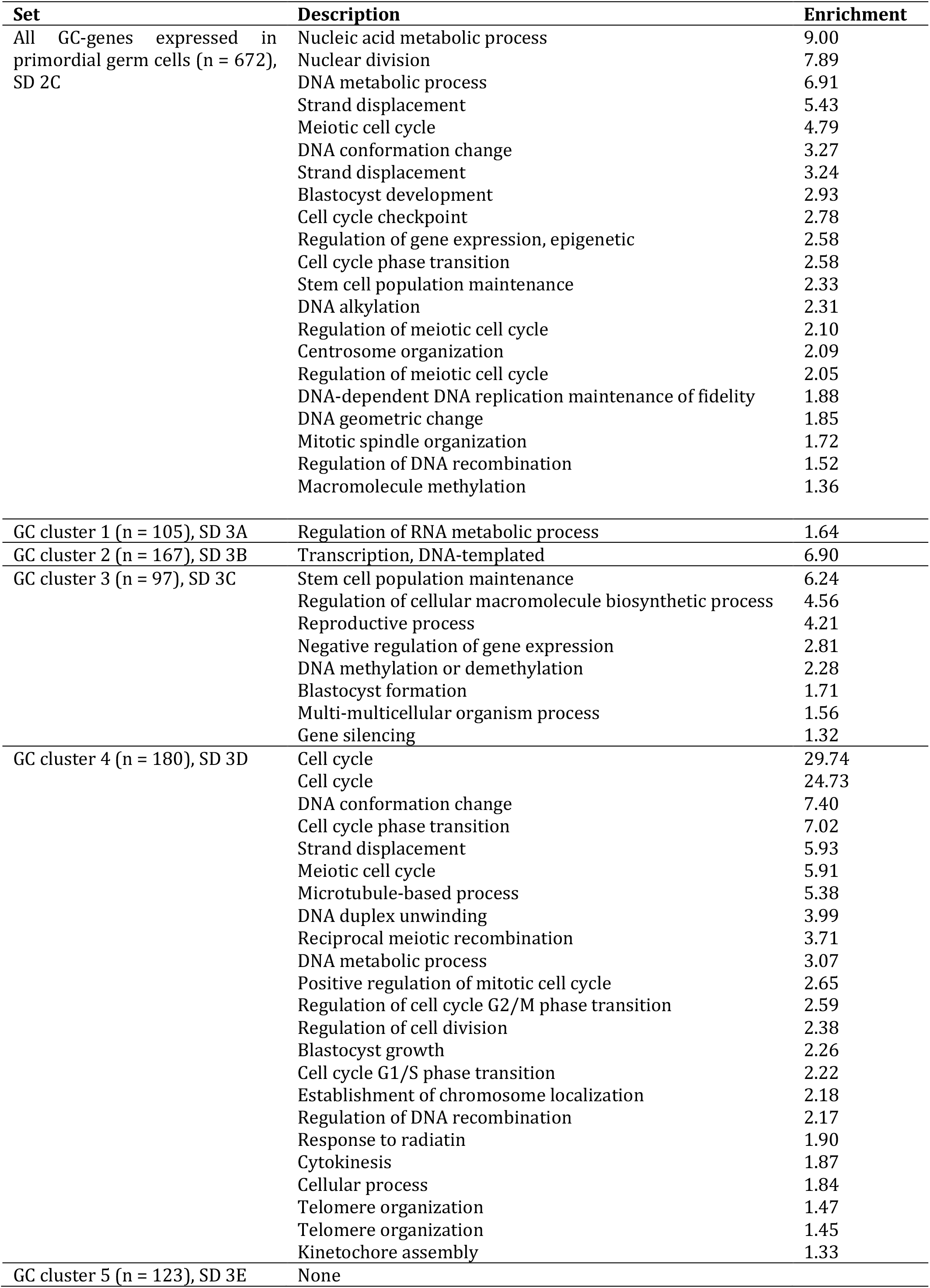
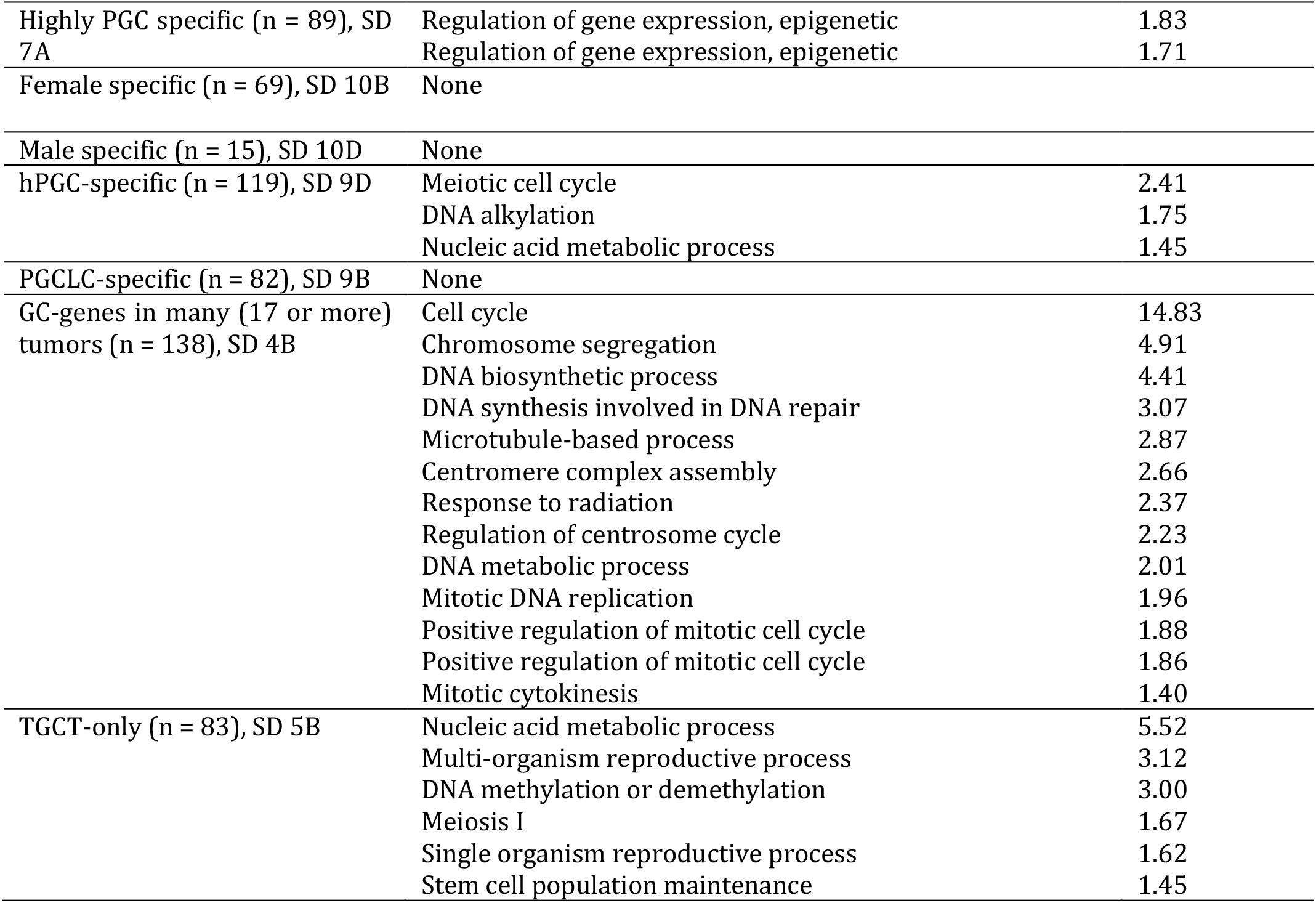
Summary of gene ontology (GO) analysis of GC-genes expressed in primordial germ cells. Enrichment equals -log_10_(p), where 1.3 is equivalent to p = 0.05 and p represents the geometric mean of p-values in an annotation cluster. Only a description of the first term of each statistically significant (enrichment > 1.3) annotation cluster is shown. Full results are shown in corresponding supplementary data (SD) for each subset.

### GC-genes can be classified in groups based on similar expression profiles in cancer

To investigate whether subgroups of embryonic GC-genes differ per tumor type, we performed an unsupervised hierarchal clustering of the 672 embryonic GC-genes and the 33 tumor types. This resulted in 5 subgroups of genes that show similar expression within tumors, and 3 subgroups of tumors that show a similar embryonic GC-gene expression profile (**Figure 2**). GC-genes in cluster 1 appeared to be particularly expressed in lower grade glioma and glioblastoma, as well as pheochromocytoma and paraganglioma, and seems to contain genes that regulate RNA metabolic processes (**Table 1 & Supplementary data 3A**). Gene cluster 2 mostly characterizes tumor group A, because it contains many genes that are expressed in acute myeloid leukemia. These genes are associated with DNA-templated transcription (**Table 1 & Supplementary data 3B**). The GC-genes in cluster 3 appeared to include the majority of genes that are highly expressed in testicular germ cell tumors and are not expressed in any other tumor type. These genes are mainly responsible for stem cell population maintenance and epigenetic changes (**Table 1 & Supplementary data 3C**). Gene cluster 4 appears to be the main determinant that separates tumor group C from A and B. Characterization of gene cluster 4 by GO analysis showed that these GC-genes are responsible for many processes related to the meiotic and mitotic cell cycle (**Table 1 & Supplementary data 3D**). A GO analysis of gene cluster 5 showed no significantly upregulated processes (**Supplementary data 3E**).

**Figure 2.**
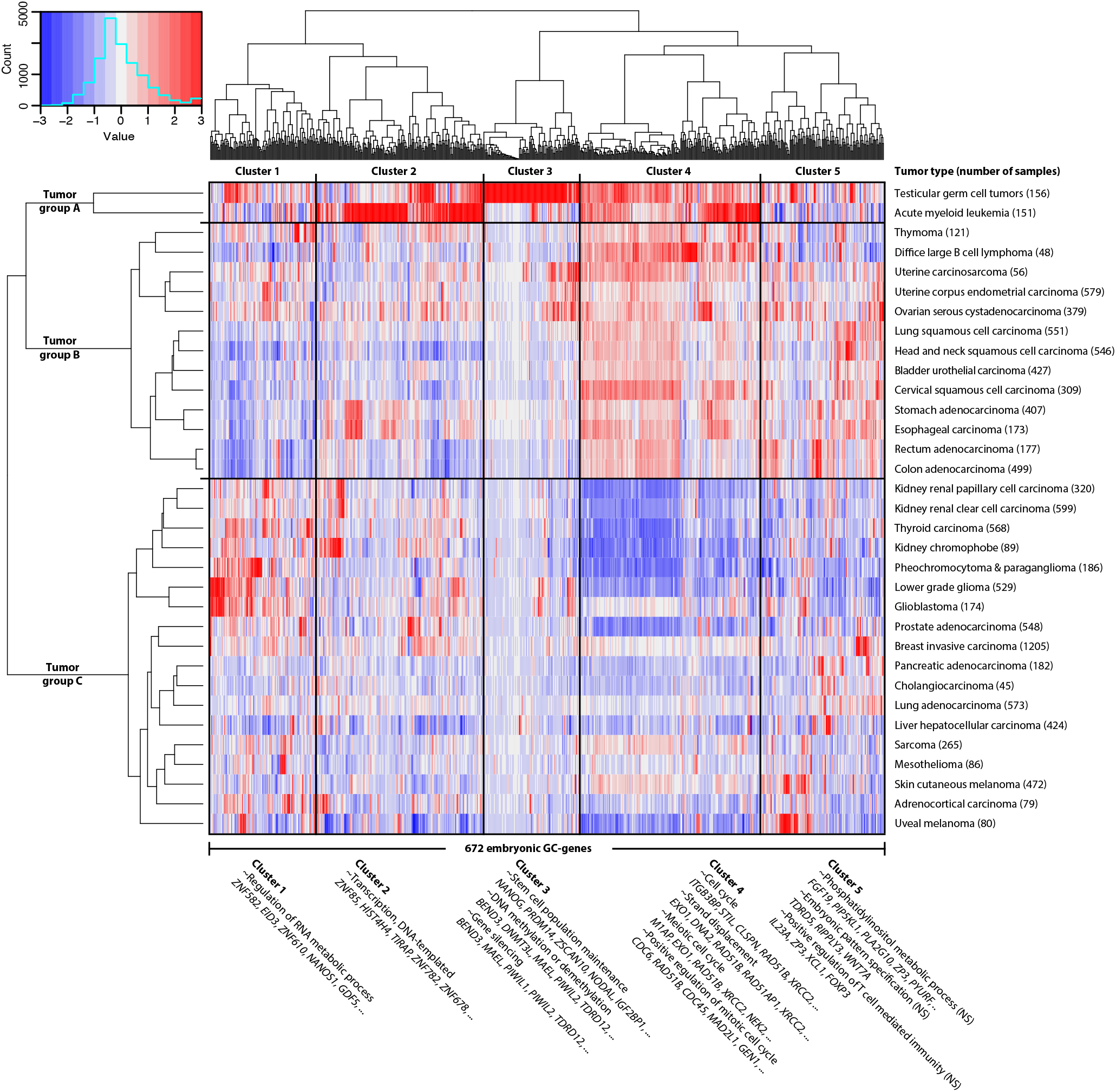
Hundreds of primordial germ cell specific genes are widely expressed in tumors. Shown here as hierarchal clustering of the average expression per tumor group (Euclidean distance, ward linkage). These 672 embryonic germline-cancer (GC) genes expressed in primordial germ cells divide tumors into two groups, mainly based on embryonic GC-gene cluster 4, which contains genes involved in the mitotic and meiotic cell cycle (table 1). Gene expression levels in tumors are indicated by a Z-score dependent color, where blue and red represent low and high expression respectively. Representative and significantly enriched GO-terms and several associated genes are shown. Cluster 5 contains no significantly (NS) enriched GO-terms.

### Embryonic GC-genes are often expressed in multiple tumor types

Besides these gene clusters, the set of 672 embryonic GC-genes expressed in PGCs contains several subgroups of interest, such as GC-genes that are expressed in more than one type of cancer. In the heat map (**Figure 2**) we observe that most genes are expressed in multiple tumor types, even though the selection criteria allow for the inclusion of genes expressed in only one tumor type. Whereas 35% of embryonic GC-genes are expressed in only one tumor type, 138 embryonic GC-genes (21%) are expressed in at least half (i.e. 17 or more) of all investigated tumor types (**Supplementary data 4A**). Due to their expression profile across tumors of different origin, we hypothesize that these GC-genes contribute to hallmarks of cancer and that tumors may be dependent on expression of a large subset of GC-genes. Characterization by a GO analysis revealed that genes expressed in 17 or more tumor types are responsible for proliferation (i.e. cell cycle processes and positive regulation of mitosis) and genome instability (i.e. chromosome segregation, DNA repair and response to radiation) (**Table 1 & Supplementary data 4B**). Opposite to this group, some genes have been included because they are expressed in only one tumor type. This particularly holds for genes expressed in testicular germ cell tumors (TGCT), as they may resemble and originate from (primordial) germ cells. 80 embryonic GC-genes (11%) have been included in our selection because of a high expression in TGCTs **(Supplementary data 5A)**, of which 70 are in gene cluster 3. Gene ontology analysis showed this subset of genes is involved in cellular aromatic compound metabolic processes, reproductive processes, DNA (de)methylation and stem cell population maintenance **(Supplementary data 5B)**. The other 592 embryonic GC-genes (89%) are expressed in a least one tumor type of somatic origin.

### A GC-gene signature score to rate shared properties between cancer and the germline

In the heat map (**Figure 2**) we observe that some tumors contain many more GC-genes than others, ranging from 84 in ovarian serous cystadenocarcinoma and head/neck squamous cell carcinoma to 360 in skin cutaneous melanoma (**Supplementary data 1C**). A tumor’s similarity to the germline thus differs vastly between tumors. In order to quantify this resemblance, we have combined our 672 embryonic GC-genes with the previously published 756 GC-genes expressed in adult male germ cells^7^ (total n = 1 143, **Supplementary data 6A**), and used the R2 bioinformatics platform^22^ to obtain a signature score for each of the 917 publicly available cancer cell lines in the Cancer Cell Line Encyclopedia^24^ (**Figure 3**). This score represents the average percentile of the GC-genes expression ranks within a particular cell line, which may be used as a measure of germ cell resemblance. Because somatic and non-somatic genes are likely to affect each other’s expression in a tumor, we also used R2 to identify key genes that are not necessarily in our dataset but correlate with the expression of GC-genes. We identified 223 genes whose individual expression positively correlates (R > 0,5 and p < 0,05) with the GC-signature scores (**Supplementary data 6B**) and 277 genes that negatively correlate (R < -0,5 and p < 0,05) with the GC-signature scores (**Supplementary data 6C**). Interestingly, only 52 of the 223 genes (23%) that positively correlate with the GC-signature are GC-genes, suggesting that genes leading to activation of the germline program in developing cancer cells may not be expressed in, or exclusive to, to the germline.

**Figure 3.**
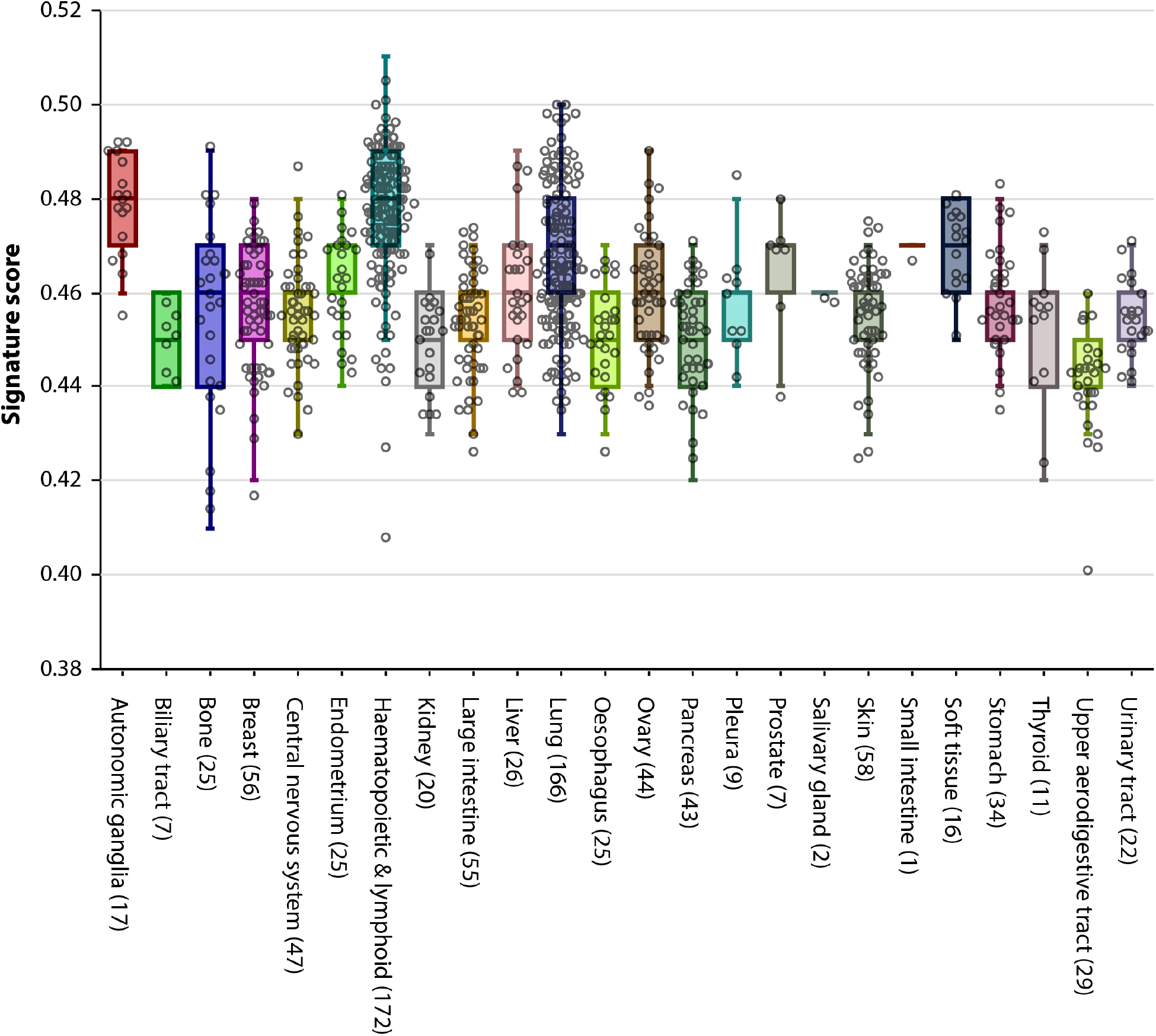
GC signature score in 917 cancer cell lines in the Cancer Cell Line Encyclopedia^24^, based on all 1 143 known GC-genes Every dot represents one cancer cell line. The signature score is the average percentile of ranked gene expression in each cell line, and may be used as a measure for a cancer cell line’s similarity to the germline.

### Expression of GC-genes is linked to increased mortality in lung adenocarcinoma

Because the lung cancer group contains sufficient samples (n = 166) and a large variability of GC-gene expression between cell lines (**Figure 3**), we used the most prevalent subtype, lung adenocarcinoma (LUAD), as a model to test whether the expression of GC-genes in this tumor-type may influence patient survival. While the decision for LUAD is based on cell line data from the CCLE, the TCGA database contains patient survival data. Using the R2 bioinformatics platform, each of the 515 patient-derived LUAD samples in the TCGA database was attributed a GC-signature score based only on the 422 GC-genes that are expressed in LUAD. Survival data shows that a high GC-gene signature score correlates with increased mortality in LUAD patients (**Figure 4**, p < 0,001).

**Figure 4.**
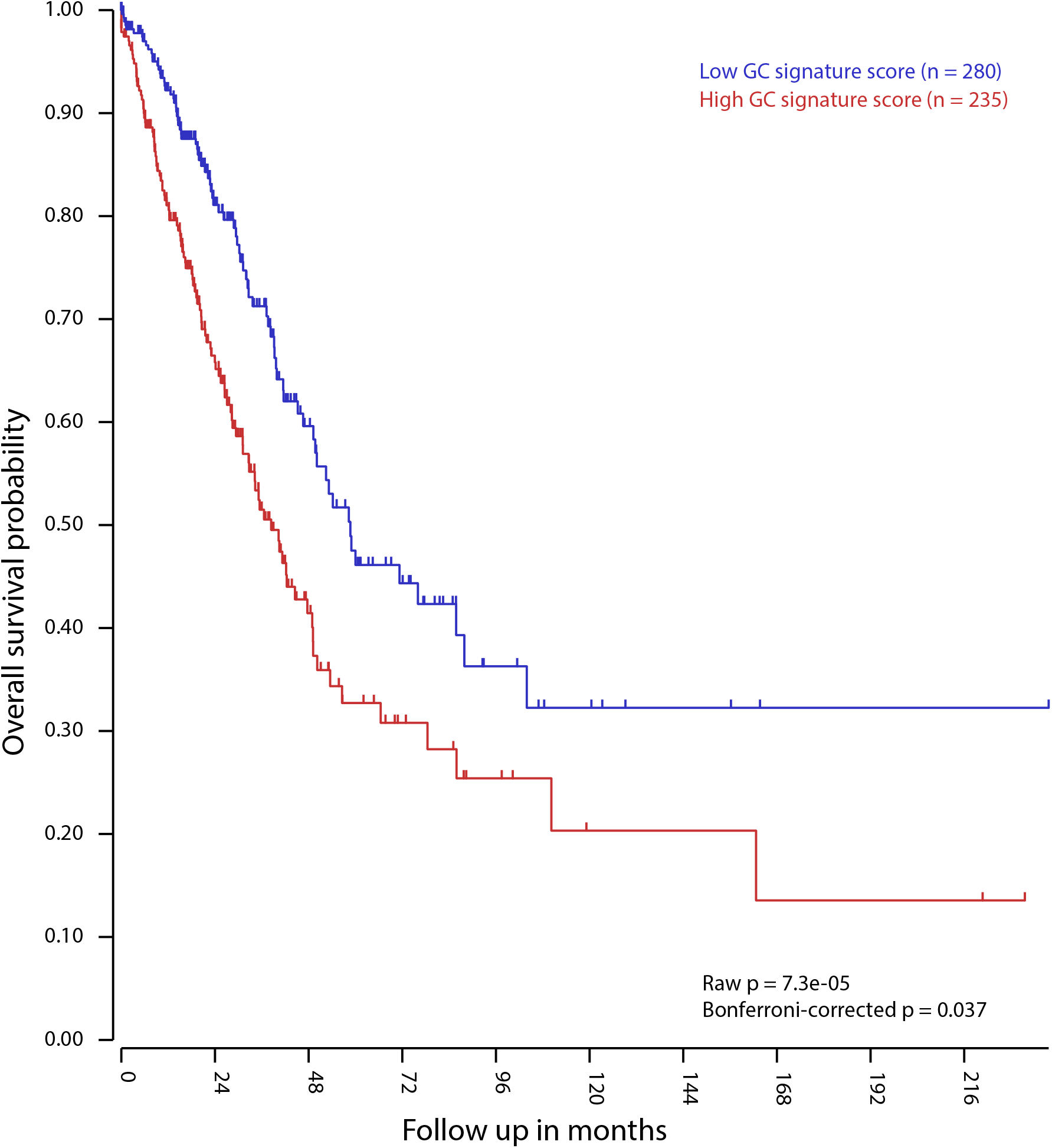
Kaplan-Meier curve of 515 lung adenocarcinoma patients, divided in two groups based on GC-gene signature score of their tumor. Figure derrived from the bioinformatics platform R2’s Kaplan Meier Scanner.

### Highly PGC-specific genes promote epigenetic alterations

We next determined which embryonic GC-genes are only expressed in ESCs and PGCs, and not in other cells of the germline. We excluded GC-genes that show expression in ovary or testis tissue in the GTEx database^18^ or adult male germ cells^17^ and required a lower expression level in somatic gonadal tissues that surround the PGCs in-situ, compared to PGCs^8^. This analysis yielded 89 embryonic GC-genes that are highly specific to the embryonic germline and cancer (**Figure 5, Supplementary data 7A**). GO analysis shows that these embryonic GC-genes are involved in regulation of epigenetic gene expression **(Table 1, Supplementary data 7B)**. Notably, 21 of these 89 embryonic GC-genes are only expressed in TGCT and not in tumors of somatic lineage.

**Figure 5.**
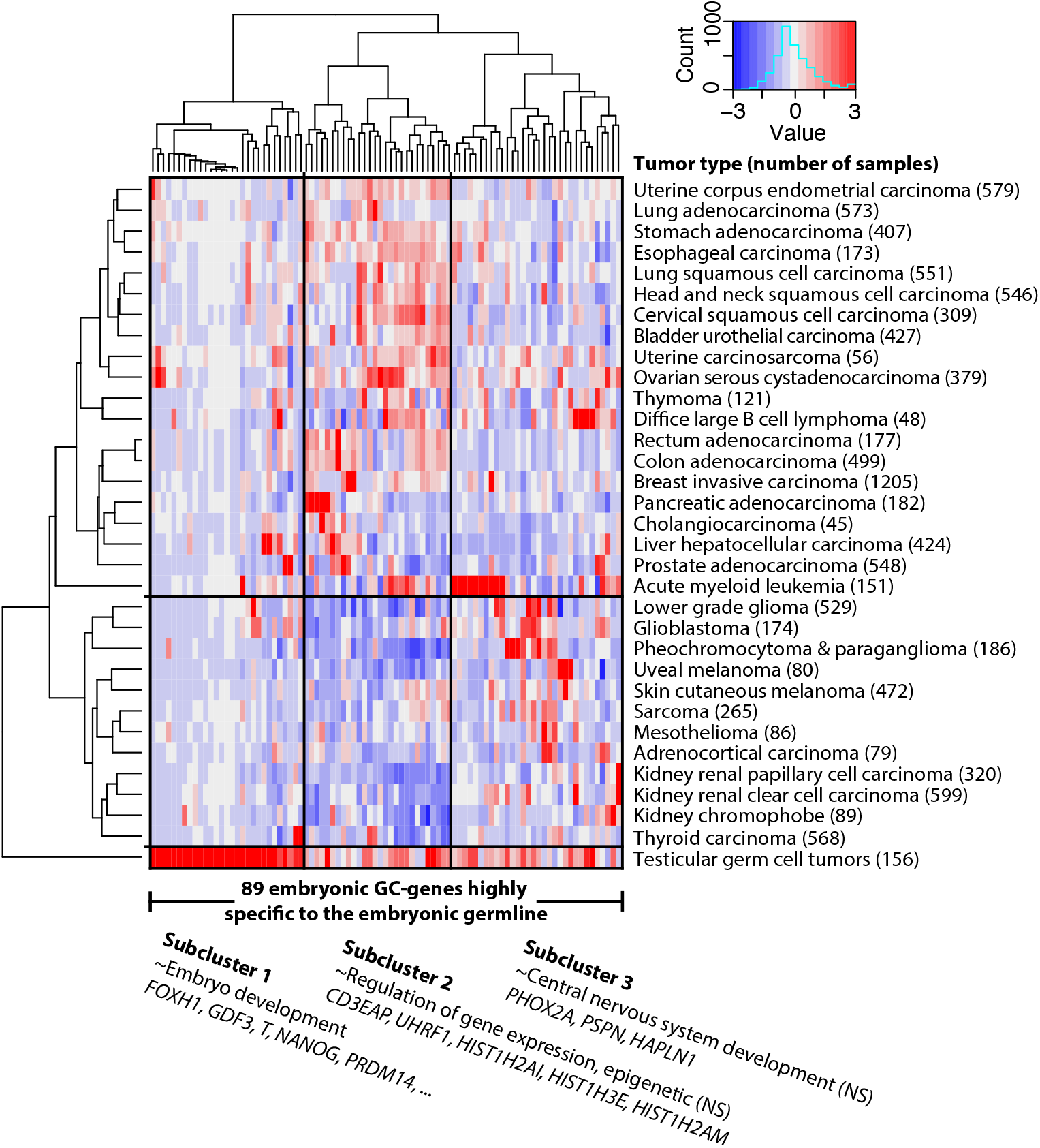
A subset of 89 embryonic GC-genes highly specific to the embryonic germline are not only expressed in germ cell tumors, but also in many tumors that originate from tissues of somatic origin. Representative GO-terms and some associated genes are known. GO-terms are not significantly (NS) enriched, possibly due to the small sample size of subclusters. Clusters in Figure 2 and subclusters in figure 5 were derrived independently.

### Cell surface molecules

Because diagnostic and therapeutic targets on the cell surface are more accessible, a final subgroup of interest are genes that encode surface proteins. We therefore used the PANTHER 10.0 classification system to analyze which embryonic GC-genes encode cell surface proteins. We identified thirteen of the 672 embryonic GC-genes (*ULBP3, GP6SPA17, CCR4, HMMR, GP1BA, KCNH5, UMODL1, WNT7A, NAT1, HYAL4, CRLF2, TNFSF4*) that are predicted to encode proteins present on the cell surface **(Supplementary data 8)**.

### Protein expression

Because RNA expression does not necessarily reflect protein expression, we compared our results to data from the human protein atlas (HPA)^23^, which contained protein expression data for 374 embryonic GC-genes (56%). By not allowing protein expression in any non-cancerous tissue other than ovary and seminiferous tubules of the testis, we identified 37 putative embryonic GC-proteins **(Table 2, Supplementary data 2A)**.

**Table 2.**
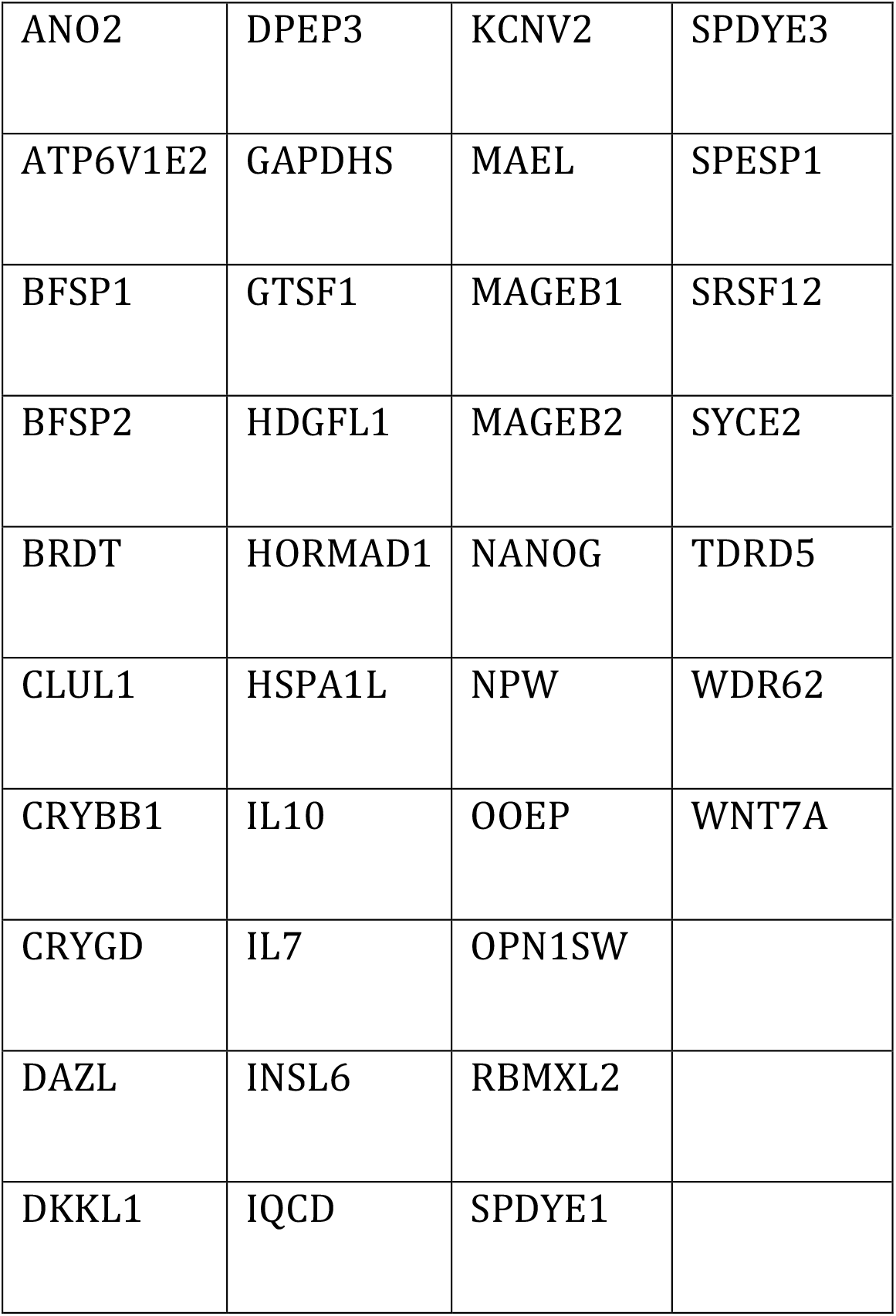
List of 37 putative embryonic GC-proteins that show no protein expression in any healthy somatic tissue, according to data from the Human Protein Atlas.

### Combination of subgroups

Finally, we searched for embryonic GC-genes that are present in multiple subgroups of interest, being (i) expression in multiple tumor types, (ii) high embryonic germline specificity, (iii) cell surface expression, and (iv) validated on the protein level (**Table 3**). We found that *NAT1* and *HYAL4* encode cell surface proteins and are highly specific to the embryonic germline. *HMMR* encodes a cell surface protein and is expressed in most (24 of 33) tumor types. *APOBEC3B, FAM111B, FAM64A, FAM86C1, SPC24, TIMM8A* and *UHRF1* are specific to the embryonic germline and are expressed in most tumor types.

**Table 3.** *GC-genes that are expressed in PGCs and fall into multiple subgroups of interest for further evaluation*.

| Gene ID | > 50% tumor types | Highly specific to embryonic germline | Cell surface | Validated on the protein level |
| --- | --- | --- | --- | --- |
| HMMR | X |  | X |  |
| NAT1 |  | X | X |  |
| HYAL4 |  | X | X |  |
| WNT7A |  |  | X | X |
| APOBEC3B | X | X |  |  |
| FAM111B | X | X |  |  |
| FAM64A | X | X |  |  |
| FAM86C1 | X | X |  |  |
| SPC24 | X | X |  |  |
| TIMM8A | X | X |  |  |
| UHRF1 | X | X |  |  |
| NPW | X |  |  | X |
| OPN1SW | X |  |  | X |
| SPDYE3 | X |  |  | X |
| BFSP2 |  | X |  | X |
| NANOG |  | X |  | X |
| CRYBB1 |  | X |  | X |

## Discussion

We here identify 672 novel germ cell cancer genes (GC-genes) that are normally expressed in human primordial germ cells but are ectopically expressed in a wide variety of tumors, most of which are tumors of somatic origin. In addition to existing GC- and cancer testis antigens (CT-genes), this expansion of the GC-gene group is of particular interest to the development of anticancer therapies, as they are not expressed in any healthy adult tissue. Of particular interest are 89 embryonic GC-genes that are not expressed in adult germ cells, whole testis tissue or embryonic somatic gonadal tissue. These genes appear involved in epigenetic regulation of gene expression and gene silencing, which is a key feature of both PGCs and carcinogenesis. Because expression of these genes is usually restricted to PGCs, and thus absent in somatic tissues and adult germ cells, targeting the genes of this subset of GC-genes could lead to fewer side effects than existing therapies.

The expression of testis^5,6,25–29^ and germ cell specific^4,7,30^ genes in tumors has been widely studied. However, gene expression of the embryonic germline had not yet been systematically compared to cancer, despite being suggested in two key publications that sparked CT-gene research^3,4^. We here show that the similarities between cancer and the embryonic germline are widespread and include processes that favor survival of the cancer cell. This finding further supports the ‘soma-to-germline’ oncogenic model, in which the upregulation of germline-specific genes promotes cancer cell development and survival through the acquisition of germ cell-like properties^31–33^ (**Figure 6**). Driven by epigenetic changes, these properties are normally strictly isolated and controlled within the germline but may allow cancer cells to prioritize their own survival over the survival of the soma. As a consequence, subsequently acquired (pseudo)meiotic functions may help the cancer cells to disturb normal cell cycle regulation and DNA repair mechanisms in order to evade checkpoints and apoptosis^32^.

**Figure 6.**
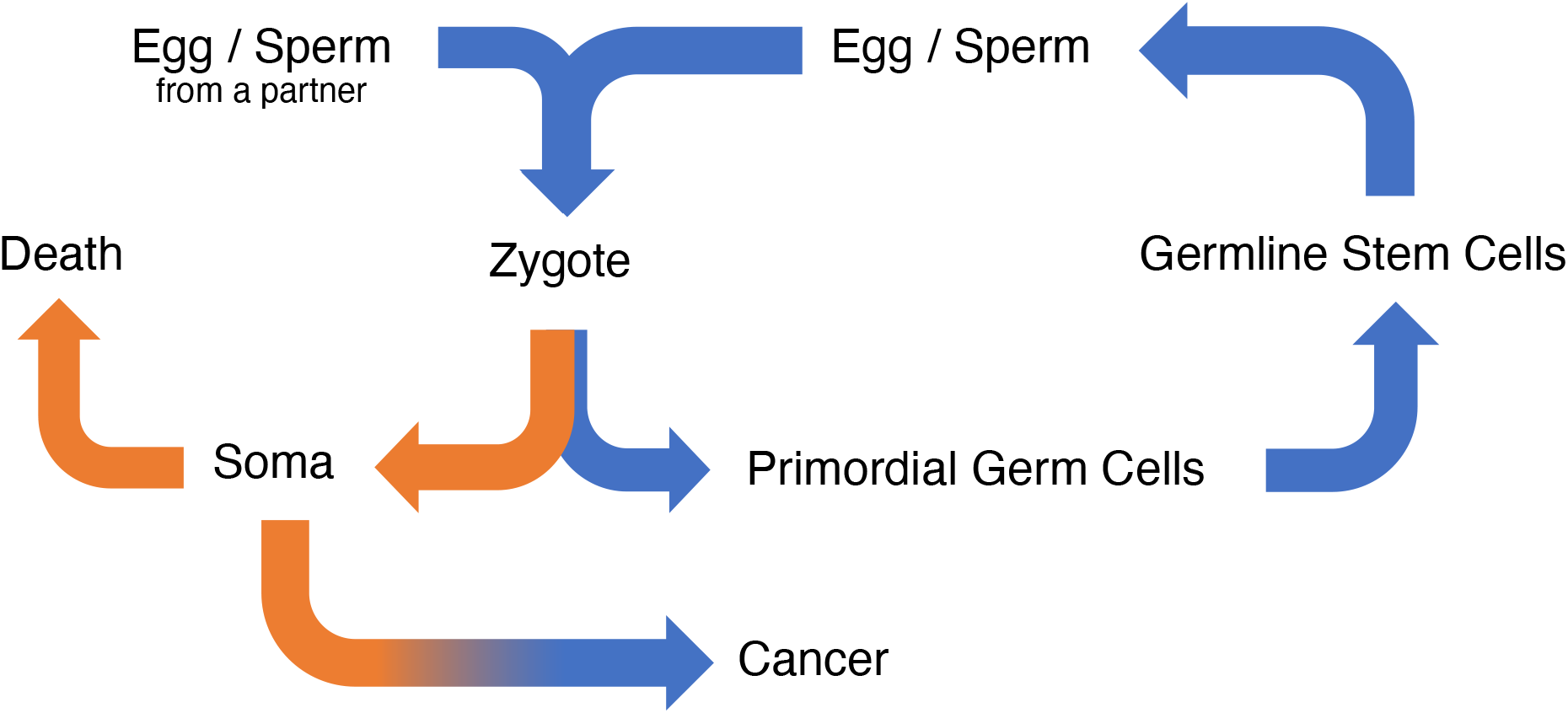
The soma-to-germline oncogenic model: processes related to the human life cycle and cancer development are closely related. Orange: somatic properties. Blue: germline properties.

Despite strict criteria for the inclusion of genes based on RNA expression, some genes may have been falsely in– or excluded. Firstly, we show that many embryonic GC-genes are involved in epigenetic alterations, a well-known mediator of oncogenesis^34^, potentially leading to the downregulation of some tumor suppressor genes. Because we selected genes based on elevated expression in tumors, downregulated (e.g. tumor suppressor) genes will have escaped our selection even though their downregulation may be specific to cancer and the germline. On the other hand, we may also have falsely included some genes, specifically those only expressed in somatic cells under specific conditions. This may for instance be the case for genes related to mitosis, as hPGCs divide mitotically before initiating meiosis in females around week 10^35^. Mitosis-associated mRNAs may therefore be relatively overexpressed in hPGCs compared to healthy somatic tissues that do not divide as rapidly. Our GO analyses show that mitosis-related processes are only enriched in cluster 4, which is the main determinant between tumor groups A/B versus C. A third reason for the false in– or exclusion of genes is the difficulty of detecting genes unique to cell types that are heterogeneous within one tissue, or expressed in other tissues than those assessed in GTEx. For example, we expect genes that are expressed in stem cells and not in differentiated tissues to be lowly expressed in the GTEx database, and thus fall below the level of exclusion. Another example is ‘whole blood’, which is one of the tissues in the GTEx database that contains many different cell types in varying numbers whose distinctions are not appreciated by our analysis. This could mean that the large number of GC-genes that is only expressed in acute myeloid leukemia, and not in any other tumor type, is not specific to AML but might also be physiologically expressed in some myeloid cells. Nevertheless, we have chosen not to exclude these genes from our analysis because they may still be relevant to the treatment development of acute myeloid leukemia, similar to why we have not excluded 83 GC-genes that have only been included due to their expression in TGCTs. Lastly, some genes with low mRNA expression show high protein expression or vice versa^36^. For example, SUV39H2 (also known as KTM1B) is expressed in nearly all tumor types. SUV39H2 is known to negatively regulate gene expression in germ cells^37^. More specifically, it is involved in cell cycle regulation, transcriptional repression and the regulation of telomere length^38,39^. Deletion of this gene allows for partial elongation of telomeres, thereby aiding tumor growth, suggesting that the presence of SUV39H2 is unlikely to promote tumor growth. This paradox may be explained by posttranscriptional regulation of gene expression, which leads to the accumulation of RNA and the absence of protein. Such posttranscriptional regulation, which also physiologically occurs in germ cell development^17^, may have led to the false inclusion of other genes as well. Because the present study is based on large datasets comprised of RNA data^8,17–19^, this remains a notable point of caution. We have attempted to validate these genes on the protein level using the HPA^23^ as a validation tool and found 37 putative GC-proteins. Despite the increasing reliability of the HPA, several drawbacks remain. Most notably, the antibodies are affinity purified and may thus not be selective to the antigen. For example, NANOG is not expressed in testis tissue according to the HPA, but a study we collaborated in has shown otherwise^40^. Therefore, future research that further explores the therapeutic relevance of individual GC-genes should also investigate the GC-gene encoded proteins or the phenotypic effects of GC-gene expression.

Because human PGCs differentiation includes dynamic events such as migration and global epigenetic resetting^14^, the stage at which they are isolated influences the results. The PGCs used in our analysis are from human embryos that are 5.5 – 8.5 weeks old, which are similar to mouse PGCs around embryonic day 13^8,41,42^. Gonadal differentiation is already initiated at this stage^14,35,42^, allowing for the attribution of meiosis-related gene activity (gene cluster 4) to the initiation of meiosis in female germ cells. While we hypothesize that PGC migration and cancer metastasis share features, genes that drive this process have probably already been downregulated in post-migratory PGCs used in this analysis. It is also unlikely to be able to observe processes related to metastasis in GC-genes expressed in PGCLCs, as they represent pre-migratory PGCs in week 2-3. While this might be due to the technical difficulty of isolating migrating PGCs, another challenge in ‘catching’ migration-related processes in our selection is that the tumor dataset only contains primary tumor samples^19^. This means that the RNAs for these genes must already be present before possible metastasis in order to be included. This could indicate that the tumors that express these genes are at an elevated risk to metastasize. Future research into the migratory potential of (early) germline cells could elucidate to what extent this is the case. In addition, further research into the expression of GC-genes in metastatic tumors may yield more putative therapeutic targets that are involved in processes that allow for metastasis.

Because human PGCs remain hypomethylated until week 16^43^, it is possible that many genes are randomly expressed in PGCs, which is a feature shared by cancer cells. While we have attempted to achieve tumor specificity through our strict inclusion criteria, a tumor’s true dependency on these genes remains questionable. Even though we have utilized GO analyses to show that embryonic GC-genes may be involved in cancer, and that this mechanism of action is plausible, some genes may have been included due to random activation as a consequence of global DNA hypomethylation. As a tumor’s dependency on a potential therapeutic target can be a requirement for the success of certain therapies^44^, this will have to be elucidated at the protein level for every individual gene level.

Hence, in addition to the previously identified CT and adult GC-genes, we here identify 672 genes that are expressed in PGCs and cancer, of which 48% has not been identified as CT or GC-gene before. Many of these genes are expressed in multiple tumor types. Because these genes are highly specific to tumors, and absent in adult germ cells and somatic tissues, targeting of their gene products is expected to lead to very limited side effects in cancer therapy. We therefore anticipate that this data will not only lead to a better understanding of tumor biology, but also to development of improved diagnostics and treatment options.

## Acknowledgements

The authors thank the De Snoo van ‘t Hoogerhuijs-stichting and the Amsterdam Research Institute Reproduction and Development for their financial support to this project.

## Author Contributions

J.W.B. and G.H. conceived and designed the study. J.W.B and J.K. performed bioinformatic analyses. J.W.B., P.L. and J.K. performed data visualization. J.W.B. and P.L. developed the GC-gene web-application. J.W.B., N.I., J.K., and G.H. interpreted the results. J.W.B., N.I., P.L., A.vP. and G.H. critically read the manuscript. J.W.B and G.H. wrote the manuscript.

---

**Supplementary figure 1.**
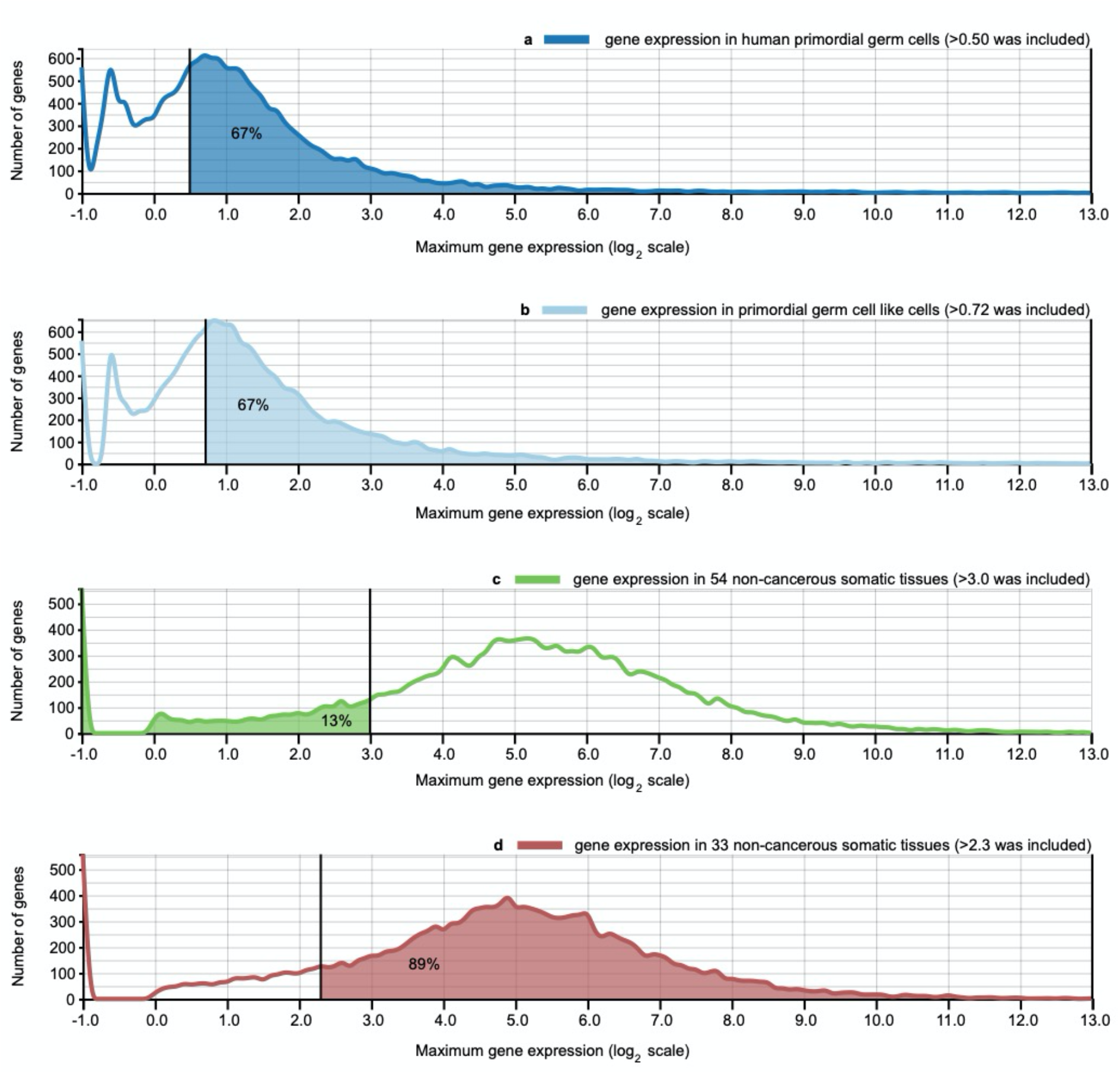
Selection of genes based on the expression in three datasets. **a**/**b**. For gene expression in human primordial germ cells, genes with a maximum gene expression in either female or male PGCs <0.72 (**a**), as well as an expression <0.50 in PGCLCs (**b**) were considered background noise and were excluded. **c**. Likewise, in order to only include genes that are exclusive to (primordial) germ cells and cancer, genes with an expression >3.0 in any normal somatic tissue were also excluded. **d**. Finally, we selected for genes with an expression >2.3 in at least one of 33 tumor types.

